# Structural analyses of the plant PRT6-UBR box in the Cys-Arg/N-degron pathway and insights into the plant submergence response

**DOI:** 10.1101/2022.08.19.504472

**Authors:** Leehyeon Kim, Chih-Cheng Lin, Ting-Jhen Lin, Yan-Cih Cao, Min-Chun Chen, Mei-Yi Chou, Wen-Hsuan Lin, Minki Kim, Jian-Li Wu, Ming-Che Shih, Hyun Kyu Song, Meng-Chiao Ho

**Affiliations:** Department of Life Sciences, Korea University, Seoul 02841, South Korea; Agricultural Biotechnology Research Center, Academia Sinica, 11529 Taipei, Taiwan; Institute of Biological Chemistry, Academia Sinica, 11529 Taipei, Taiwan; Institute of Biochemical Sciences, National Taiwan University, 10617 Taipei, Taiwan

**Author notes:** These authors contributed equally (Leehyeon Kim, Chih-Cheng Lin).

## Abstract

The submergence response in higher plants is highly dependent on the protein stability of group VII ethylene response factors, which are primarily degraded through the oxygen-dependent Cys-Arg branch of the N-degron pathway of targeted proteolysis. Knockout of PRT6, an E3 ligase and a vital component of the N-degron pathway, improves submergence tolerance in *Arabidopsis* and barley but is associated with side effects such as germination deficiency. In this study, we determined structures of rice and *Arabidopsis* PRT6-UBR box in complex with various Arg/N-degron related peptides. We identified two highly conserved motifs in the plant PRT6-UBR box, which is responsible for Cys-Arg/N-degron recognition. Structural and mutagenesis studies revealed the importance of two conserved motifs for Cys-Arg/N-degron recognition. The phenotype of *Arabidopsis* seedlings with *PRT6-UBR* mutants in these newly identified conserved motifs showed superior submergence survival suggesting that rational manipulation of the PRT6-UBR box can improve flood tolerance. Our results provide an engineering platform for generating crops with improved submergence tolerance.

## Introduction

Although oxygen produced by plants is the major source released into the atmosphere, oxygen is also essential for plant for respiration to generate energy and to conduct some cellular functions. Plant submergence limits oxygen exchange, resulting in cellular oxygen reduction, in turn threatening survival. In many places around the world, climate change has transformed once-in-a-lifetime heavy rainfall into frequent occurrences thus increasing the likelihood of submergence and, along with it, devastating agricultural economic loss and global food security risk^1^. Accordingly, it is critical to develop strategies to improve plant submergence tolerance.

Submergence is a complex stress that affects hormone signalling, carbohydrate metabolism, redox balance, cytosolic pH, reactive oxygen species (ROS) level, and other plant physiology^2^. Recent studies have shown a direct association between submergence and the group VII ethylene response factors (ERF-VIIs). The expression and degradation of ERF-VII proteins is highly correlated with plant submergence survival^3–10^. One of the characteristic features of ERF-VIIs is their conserved N-terminal domain, which possesses Met-Cys-Gly-Gly as the first four amino acid residues. This conserved N-terminal MCGG motif is the N-degron for the oxygen-dependent Cys-Arg/N-degron pathway of targeted proteolysis, which leads to destabilization of ERF-VIIs the presence of oxygen. As the initial step in this pathway, the first methionine is removed by methionine aminopeptidase, exposing the cysteine residue at the N-terminus^11^. This N-terminal cysteine is then oxidized in the presence of oxygen by plant cysteine oxidases (PCOs) converting it into a negatively charged cysteine sulfonate (CysO_2_); subsequent arginylation by the arginyl-tRNA protein transferase 1 (ATE1) creates the unique Cys-Arg/N-degron^12,13^. This N-degron possesses a positively charged arginine and a negatively charged cysteine sulfinate at the first and second positions, respectively. The N-terminal Arg residue is the primary destabilizing residue in the Arg/N-degron pathway^12^ and, in this report, we use the term ‘secondary destabilizing residue’ to refer to the second residue of the Arg/N-degron (cystiene sulfinate). Once formed, the Cys-Arg/N-degron is recognized by the UBR box of PRT6, an E3 ubiquitin ligase for polyubiquitylation, resulting in targeted proteolysis by the ubiquitin–proteasome system (UPS)^12,14,15^.

Overexpression of *Arabidopsis* ERF-VIIs often upregulates downstream genes for hypoxia responses and improves tolerance to submergence, suggesting that higher levels of ERF-VIIs enhance plant submergence survival^4^. Indeed, endogenous ERF-VIIs accumulation in barley, achieved by lowering PRT6 levels to impede the Arg/N-degron pathway, improves tolerance to waterlogging^4,11^. Similarly, *prt6* knock-out in *Arabidopsis* enhances tolerance to submergence^16^. The *prt6* mutant allele, *ged1*, which possesses a non-functional PRT6 protein, shows high tolerance to starvation during submergence or prolonged darkness^17^. Knockout of *prt6* in barley and *Arabidopsis* also enhance tolerance to other abiotic stresses, such as salinity and drought^11,15^. All these findings suggest the possibility of introducing non-functional *prt6* mutants into crops for enhanced tolerance to several abiotic stresses. However, *prt6* knockout lines are also associated with disadvantages^15^. For example, *prt6* mutants show defects in the development of shoots and leaves, and are hypersensitive to abscisic acid (ABA), resulting in slower germination and lipid breakdown^18,19^. A recent study suggested that PRT6 activities are required to optimize seed storage reserves to coordinate germination and seedling establishment^20^. In short, while developing flood-resistant crops based on PRT6 modulation may be a promising strategy to improve submergence tolerance, careful manipulation of PRT6 activities is required to avoid unwanted effects.

Herein, we report the crystal structures of the UBR boxes of *Oryza sativa* (rice) PRT6 and *Arabidopsis thaliana* PRT6 (*Os*PRT6-UBR and *At*PRT6-UBR, respectively) in complex with different Arg/N-degron related peptides at 1.5–2.4 Å resolution. Our sequence and structural studies showed that two highly conserved motifs near the Arg/N-degron binding pocket are critical for recognition of the oxidized cysteine (CysO_2_). Mutagenesis of these conserved motifs of *Os*PRT6-UBR led to a dramatic reduction of binding affinity to the Cys-Arg/N-degron mimic, an Arg–Asp-containing peptide. Furthermore, overexpression of these PRT6-UBR mutants in *prt6*-knockout *Arabidopsis* seedlings showed much better submergence tolerance compared to those overexpressing wild-type PRT6-UBR. Our study thus demonstrates that structure-guided modification of PRT6-UBR provides a possibility to improve submergence tolerance in crops.

## Results

### Motifs that are highly conserved in plant PRT6-UBR box

Phylogenetic analyses of UBR box motifs from plant PRT6s, *S. cerevisiae* UBR1, and animal UBRs showed that they all share a common ancestor, but evolved into three distinct clades (Supplementary Fig. 1), a finding that is consistent with a previous report^21^. Amino acid sequence alignment showed that the residues to form a three-zinc-stabilized heart-shaped fold^22^ and the Nt-Arginine of Arg/N-degron binding are highly conserved, except for one C-terminal histidine in plants and animals that holds structural zinc; in yeast an N-terminal histidine fulfills the same purpose. Based on the reported structures of yeast and human UBR boxes^23,24^, residues for Arg/N-degron second destabilizing residue recognition can be predicted. Of note, surprisingly, our phylogenetic analyses showed that two motifs near the Arg/N-degron second destabilizing residue binding site are only conserved within plant PRT6, suggesting its role in CysO_2_ recognition (Fig. 1 and Supplementary Fig. 1).

**Figure 1.**
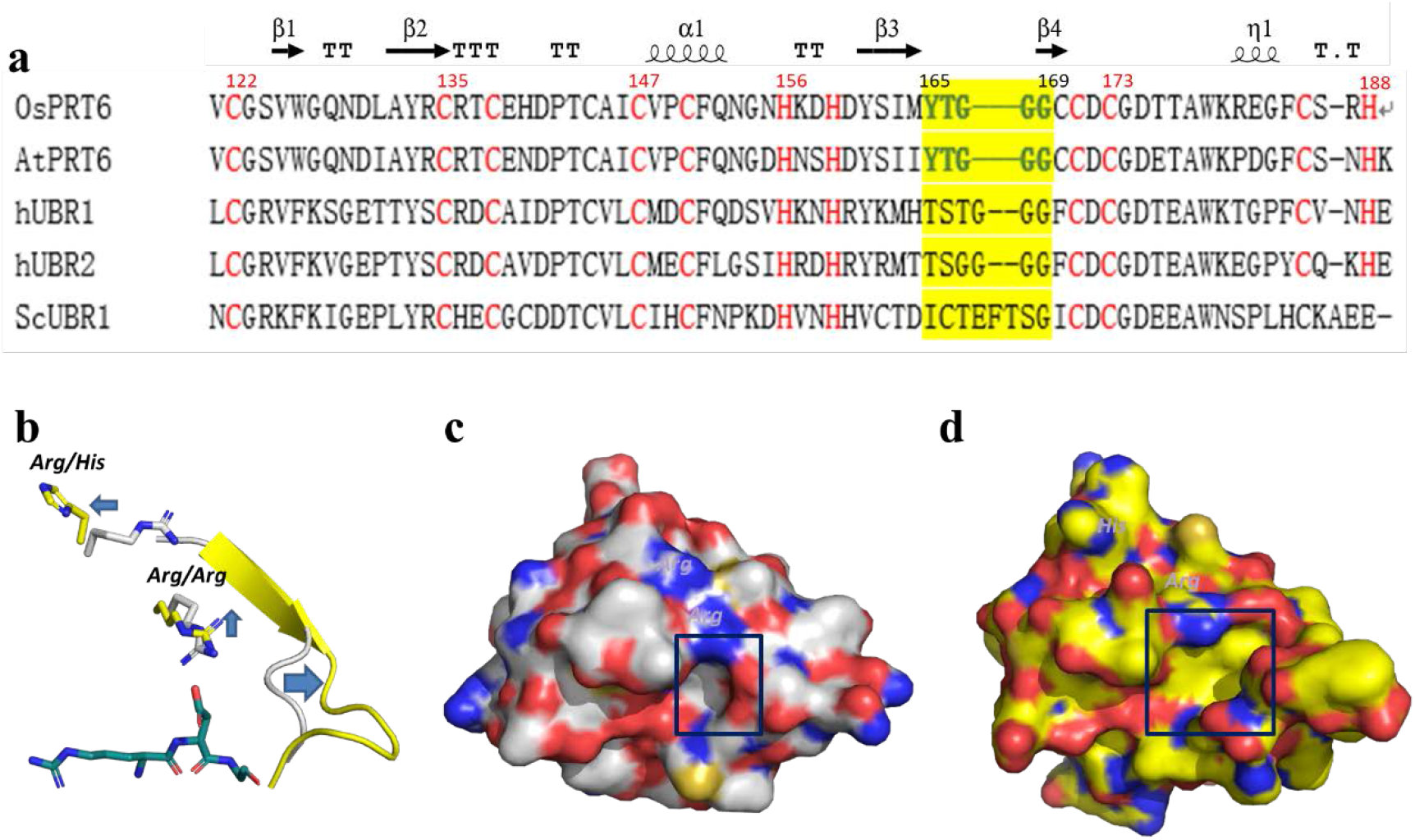
Structural features of plant PRT6-UBR box. (a) Structure-based sequence alignment with secondary structure was calculated by PROMALS3D using *Os*PRT6-UBR (this study. PDB ID: 7KUW), *At*PRT6-UBR (this study, PDB ID: 7XWD), *Hs*UBR1-UBR (PDB ID: 3NY1), *Hs*UBR2-UBR (PDB ID: 5TDA) and *Sc*UBR1-UBR (PDB ID: 3NIH). The yellow highlight indicates the structurally most divergent region among the various UBR boxes. (b) Overlay of the Tyr165–Gly169 region and nearby key Arg residues (Arg134/Arg136 in *Os*PRT6-UBR; Arg136/His138 of *Sc*UBR1-UBR). The *Os*PRT6-UBR and *Sc*UBR1-UBR are shown as a cartoon diagram in gray and yellow, respectively. The bound N-degron peptide (Arg-Asp-Ala-Ala) in *Sc*UBR1-UBR is shown as a green stick to indicate the binding location of the Arg/N-degron secondary destabilizing residue. The arrows show the structural deviations between *Os*PRT6-UBR and *Sc*UBR1-UBR. (c, d) Surface map of *Os*PRT6-UBR and *Sc*UBR1-UBR boxes are shown. Color scheme: red, negatively charged residues; blue, positively charged residues; gray (*Os*PRT6-UBR)/yellow(*Sc*UBR1-UBR), neutral. The binding pocket for Arg/N-degron secondary destabilizing residues are shown by the black boxes.

### Unique structural features of *Os*PRT6-UBR

To further investigate the structural uniqueness near the secondary destabilizing (CysO_2_) binding site of the UBR box in plants, we determined the structure of the *Os*PRT6-UBR at 1.6 Å resolution. *Os*PRT6-UBR shares 38 and 36% amino acid sequence identity, and 54 and 46% sequence similarity with *Hs*UBR1-UBR and *Sc*UBR1-UBR, respectively. *Os*PRT6-UBR structurally resembles *Hs*UBR1-UBR and *Sc*UBR1-UBR with Cα root-mean-square deviations (RMSD) of 1.5 and 1.8 Å, respectively^23,24^. Therefore, the folding pattern, secondary structural elements, and zinc-coordinating residues of *Os*PRT6-UBR are virtually the same as the human and yeast UBR box; however, there is significant structural deviation in the loop region (Tyr165–Gly169), which is only conserved among plant PRT6-UBRs (Fig. 1a). This loop is right after the β3 strand (Tyr160–Met164) and adopts a 90 degree turn to connect to the highly conserved Asp172, which interacts with the positively charged α-amino group of the N-degron. The Tyr165–Gly169 loop, which is one residue shorter than that of *Hs*UBR1-UBR and three residues shorter than that of *Sc*UBR1-UBR, forms part of the binding pocket for the Arg/N-degron secondary destabilizing residue (Fig. 1a and Supplementary Fig. 1). Because the His156 and the Cys171 are coordinating a zinc atom, the positon of the β3 strand and key recognizing Asp172 residue are fixed. Therefore, the shorter Try165–Gly169 loop forms a sharp turn (Fig. 1b) resulting in a narrower Arg/N-degron secondary destabilizing residue binding pocket (7.8 Å in length and 6.4 Å in width) as compared to the *Sc*UBR1-UBR (PDB ID: 3NIL), which has modest-sized pocket (12.2 Å in length and 6.8 Å in width) and *Hs*UBR1-UBR (PDB ID: 3NY1), which has the largest pocket (14.4 Å in length and 10.2 Å in width) (Fig. 1c, d).

The other difference is the conserved Arg134–Glu139 sequence (Arg-Cys-Arg-Thr-Cys-Glu). Cys135 and Cys138 are part of the C2H2 zinc finger and thus are conserved in all UBR boxes but the unique salt bridge between Arg136 and Glu139 is only found in plant PRT6-UBR. The Cys135 carboxyl oxygen forms a 4-residue β-hairpin with the Asp139 amide, forcing both positively-charged side chains of Arg134 and Arg136 to face the same side. Therefore, the Arg134 side chain points toward the Arg/N-degron secondary destabilizing residue (CysO_2_) binding pocket (Fig. 1b)

### Structure of *Os*PRT6-UBR in complex with N-degron peptides

The crystal structure of *Os*PRT6-UBR in complex with a tripeptide, Arg-Asp-Gly, a mimic of the Cys-Arg/N-degron physiological substrate^25,26^, was determined at 1.6 Å resolution (Fig. 2a and Supplementary Table 1). The arginine, the Arg/N-degron primary destabilizing residue, is surrounded by a negatively charged environment that includes Asp141, Asp172 and Asp175, which are highly conserved in all UBR boxes (Fig. 1a and Supplementary Fig. 1). This arginine also interacts with Asp141 and Thr177 via water molecules. The backbone of the N-degron packs tightly into the groove by forming three coordinating hydrogen bonds with the backbone carbonyl group of Cys170 and the backbone amide nitrogen of Cys170 and Thr143 (Fig. 2b and 3a). The aspartate residue at the second position residue of Arg/N-degron forms a salt bridge with Arg134 at the distance of 2.8 Å. When the aspartate, the Arg/N-degron secondary destabilizing residue, is replaced with serine, which possesses a shorter and polar side chain, the Arg134 and serine can also form a hydrogen bond with a distance of 3.1 Å (Figs. 2c and 3d).

**Figure 2.**
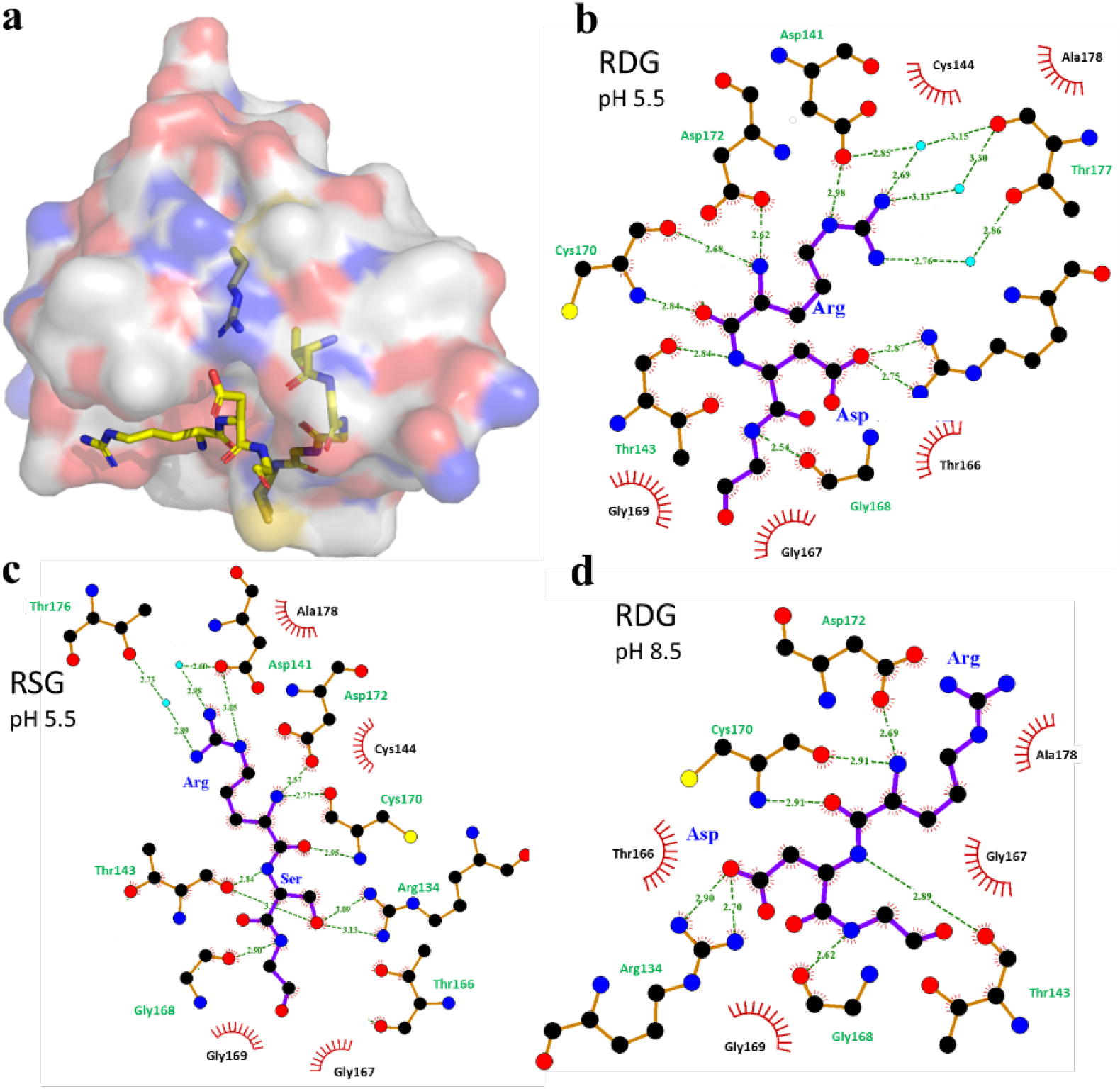
The interaction between *Os*PRT6-UBR and Arg/N-degron peptides. (a) The *Os*PRT6-UBR is drawn as a surface presentation. The key interacting residues (Arg134 and Tyr165-Gly169) and RDG peptide are shown as a yellow stick model. (b-d) The RDG-*Os*PRT6 at pH 5.5 (b), RSG-*Os*PRT6 at pH 5.5 (c), and RDG-*Os*PRT6 at pH 8.5 (d) interaction diagrams are displayed by LIGPLOT+^51^. The N-degron peptides and *Os*PRT6 residues are shown as purple and orange lines. Bound N-degron peptides and *Os*PRT6-UBR protein residues are labeled in blue and green colors, respectively. The water molecules are shown as cyan balls. The hydrogen bonds are shown as green dashed lines and the spoked arcs represent van der Waals interactions.

**Figure 3.**
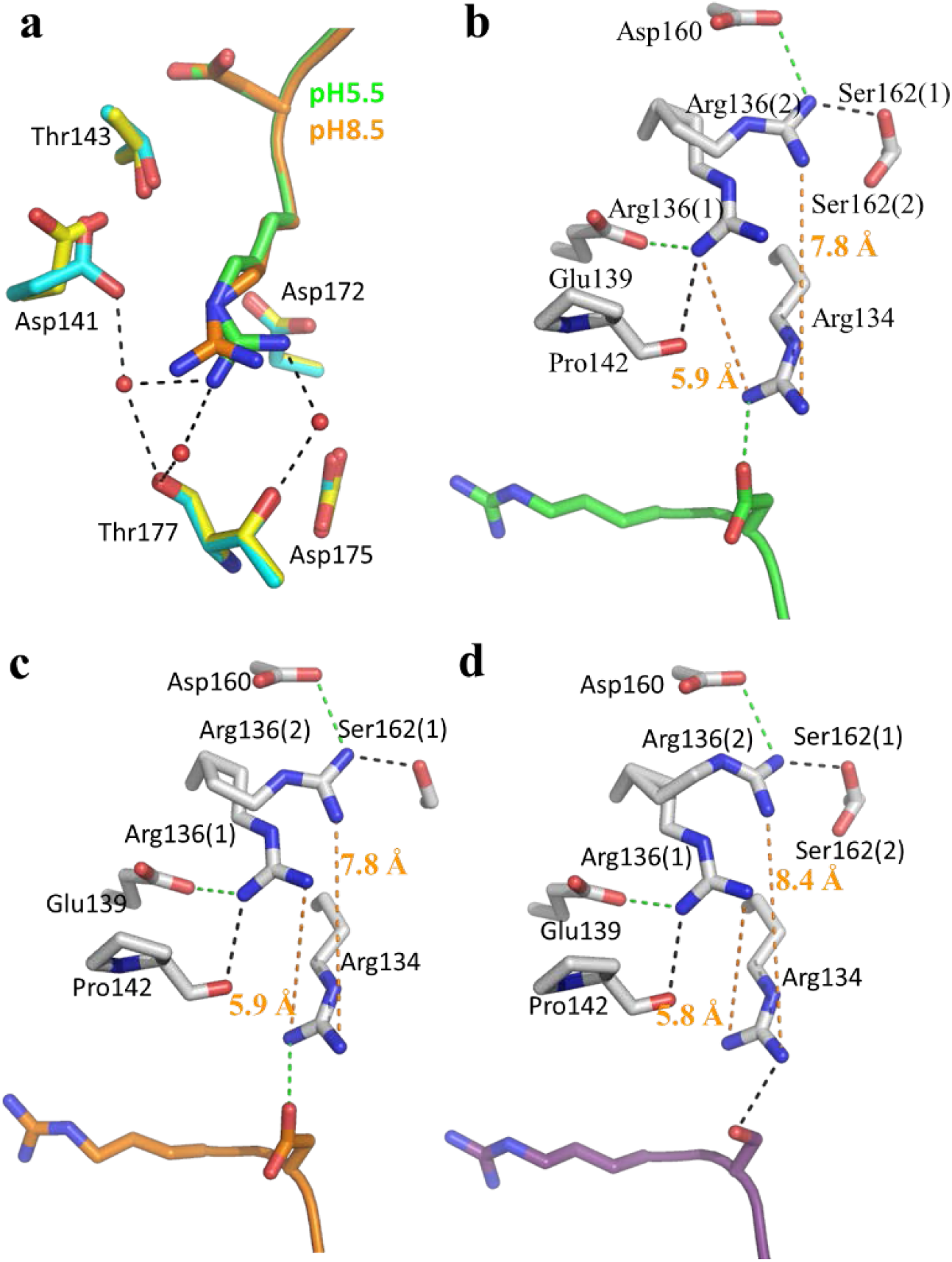
The dual conformations of two Arg residues of *Os*PRT6-UBR with various Arg/N-degron peptides. (a) The Arg of Arg/N-degron conformation changes upon pH change. The *Os*PRT6-UBR and bound RDG peptide at pH 5.5 are shown as cyan and green sticks, respectively, and at pH 8.5, they are shown as yellow sticks and orange sticks, respectively. The water molecules are shown as red balls. (b) The conformations of Arg136 and Arg134 in *Os*PRT6-UBR/RDG complex at pH 5.5. (c) The conformations of Arg136 and Arg134 in *Os*PRT6-UBR/RDG complex at pH 8.5. (d) The conformations of Arg136 and Arg134 in *Os*PRT6-UBR/RSG complex at pH 5.5. (b-d) *Os*PRT6-UBR is shown as gray sticks. The bound RDG peptide at pH 5.5 and pH 8.5, and the bound RSG peptide at pH 5.5 are shown as green, orange, and purple sticks respectively. Ser162 side chain in *Os*PRT6-UBR also has dual conformations. The hydrogen bond and charge-charge interactions are indicated as black and green dashed lines. The orange dashed line indicates key distance. Key residues are labeled in black.

During hypoxia, the cytosolic environment changes from neutral to acidic^27–29^ and the recombinant UBR box binds to the Arg/N-degron better at acidic pH^23,30^. Therefore, we determined *Os*PRT6-UBR structures in complex with the RDG peptide at pH 5.5 and pH 8.5. Compared to pH 8.5, at pH 5.5, we observed that the arginine side chain of the N-degron moves closer to Asp141, by 1.2 Å, to form an additional hydrogen bond network via water molecules (Figs. 2b, 2d and 3a).

We also noticed two alternative conformations of the Arg136 side chain (Fig. 3b-d). In one conformation, the Arg136 side chain, tilting away from Arg134, forms a salt bridge with the Asp160 side chain and a hydrogen bond with the Ser162 hydroxyl group (Fig. 3b-d). In the other conformation, the Arg136 side chain tends to form a salt bridge with Glu141 and hydrogen bonds with the carbonyl backbone of Pro142, moving closer to Arg134 from a distance of 7.8 to 5.9 Å. Based on the electron density and side-chain occupancy, we observed that the Arg136 side chain prefers to move toward to Arg134 at pH 5.5. This long range charge–charge repulsion can be strong even at distances of 5–10 Å^31^. This Arg136 side chain movement towards to Arg134 results in the Arg134 side chain entering the binding pocket of the Arg/N-degron secondary destabilizing residue, which may enhance the binding of *Os*PRT6-UBR to the negatively-charged Arg-CysO_2_ residue.

### Recognition of the second position of the Cys-Arg/N-degron by *Os*PRT6-UBR

To examine the preference for the secondary destabilizing residue – the residue right after the N-terminal Arg of *Os*PRT6-UBR – we measured the binding affinities of RXG where X can be Arg, Asp, Ser and Leu as representatives of residues with positively charged, negatively charged, polar, or hydrophobic side-chains, respectively. As described above, Asp can mimic CysO_2_^25,26^. Geometries of both the sulfinate (–SO_2_) and carboxylate (–CO_2_) groups are approximately trigonal planar, and the distance of the S–O bond is 0.1 Å longer than a C–O bond. The atomic radius of sulfur is 0.2 Å larger than that of carbon. The pKa values of cysteine sulfinic acid (CysO_2_H) and carboxylic acid of Asp (CO_2_H) are ~2.0 and 3.9, respectively^32^. Based on geometry and pKa, the negatively charged Cys-sulfinate could have a stronger interaction with positively charged residue(s) of *Os*PRT6-UBR compared to the carboxylate side chain of Asp as a mimic of Cys-Arg/N-degron in this study.

To our surprise, rice PRT6-UBR prefers Leu as secondary destabilizing residue and shows relatively weak preference for Asp at this position (Fig. 4a). The weak binding must be still strong enough to be recognized and to allow the protein to undergo subsequent ubiquitination and degradation. To further dissect the specific determinants for the recognition of CysO_2_ by plant PRT6-UBR, we determined the crystal structures of *At*PRT6-UBR in complex with RR- and RL-peptides, two representative Arg/N-degrons (Supplementary Table 2 and Supplementary Fig. 2). Our structural analyses showed that the movement of key active site residues, in particular *At*PRT6-UBR Arg133 (corresponding to *Os*PRT6-UBR Arg134), creates space to accomodate different secondary destabilizing residues, which is similar to yeast UBR1^23^.

**Figure 4.**
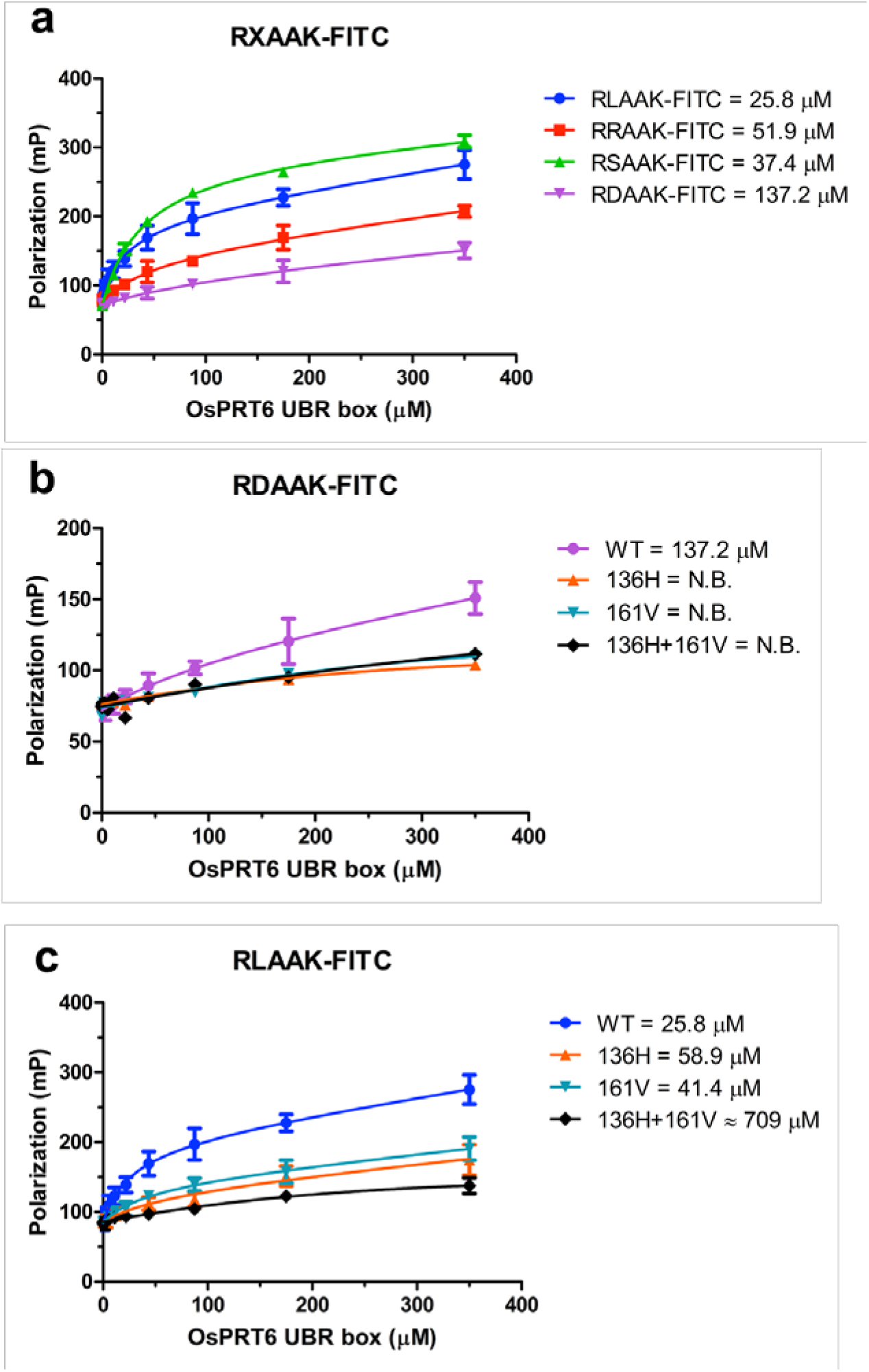
The binding affinity of measurement between *Os*PRT6-UBR proteins and Cys-Arg/N-degron peptides. (a) Measurements of fluorescence polarization of FITC-labeled RX-peptide with increasing concentrations of MBP-*Os*PRT6-UBR proteins. Measurements of fluorescence polarization of FITC-labeled RDAAK-peptide (b) and RLAAK-peptide (c) with increasing amount of wild-type, 136H, 161V, or 136H+161V mutants. For a clear signal, His-MBP-*Os*PRT6-UBR protein was used. In contrast to the wild-type protein, mutants at the putative determinant regions possess no binding affinity (N.B.) when RDAAK, a mimic of the RCO_2_-peptide, was used. The RL-peptide did not show significant discrepancy among the proteins. The error bars represent standard error of the mean of more than three independent experiments.

### Two plant-conserved motifs for Arg-CysO_2_ recognition

To reveal the role of the conserved motifs we had identified, residues 136–139 (R^136^-T-C-E^139^) and 161–168 (Y^161^SIMYT--GG^168^) were replaced by the corresponding residues from *Sc*UBR1-UBR: HECG (named 136H) and VCTDIDCTEFTS (named 161V), respectively. In addition, a double mutant, containing both 136H and 161V, was generated (136H+161V). The binding affinity between wild-type *Os*PRT6-UBR and RD-peptides, a Arg-CysO_2_ mimic was approximately 137.2 μM (Fig. 4a). However, all three mutants showed a drastic decrease in binding affinity. There was essentially no detection of binding affinity using fluorescence polarization methods. In contrast, the RL-peptide bound to the wild type protein with 25.8 μM affinity (Fig. 4b). Although the 136H+161V double mutant shows reduced binding with Kd of 709 μM, the 136H and 161V mutants showed Kds of 58.9 and 41.4 μM, respectively, which are slightly lower than the wild type protein (Fig. 4b). This clearly shows that the conserved mutated region surrounding the negatively charged residue at the second position in plants is particularly sensitive to any variation.

It is worth mentioning that despite there being weak detection of binding of *Os*PRT6-UBR to RD peptide *in vitro* and no detect of binding of OsPRT6-UBR mutants to RD peptide *in vitro*, we did see *ex vivo* activity (see next section), suggesting that even extremely weak binding can generate a physiological consequence. Alternatively, the real substrate, Arg-CysO_2_-protein, may have higher binding affinity to the wild-type and mutant PRT6-UBR proteins.

### Two plant-specific motifs for submergence survival in *Arabidopsis*

Knockout of *prt6* can significantly enhance submergence survival by inhibiting the Arg/N-degron pathway^17^. To determine whether the highly conserved motifs in plant PRT6-UBR boxes are important for submergence survival (Fig. 1a and Supplementary Fig. 1), different 14-day seedlings, including Columbia-0 (Col-0), *prt6-5* (*prt6* knockout)^33^, and *prt6-5* knockout with overexpressed *Os*PRT6 variants, were subjected to 32–33 hours of dark submergence followed by four days of recovery. We determined the damage index based on the ratio of chlorotic leaves (over 1/3 of leaf area) to overall leaves for each seedling^34^. For Col-0, only 10% of seedlings had minor damage (less than 25% damage), but 40% of seedlings had severe damage (more than 50% damage). On ther other hand, whereas the *prt6-5* line (*prt6* knock-out) showed better submergence tolerance with 45% of seedlings showing minor damage and less than 20% of seedlings showing severe damage, which is consistent with previous reports^16^. Previously, we identified *Os*ERF66 and *Os*ERF67, the rice ERF-VIIs involved in the SUB1A-1 regulatory cascade^35^. When *Os*ERF66 is overexpressed in the *prt6-5* line, we saw no significant improvement in the damage index compared with *prt6-5* line. Thus, overexpression of *Os*ERF66 in the *prt6-5* line *(prt6-5/ERF66* line) had a negligible effect in our rescue experiment. Overexpressing wild-type *Os*PRT6 in the *prt6-5/ERF66* lines (three different lines) showed a similar damage index to Col-0 (Fig. 5). In short, *Os*PRT6 can rescue *At*PRT6 function in *prt6* line as a positive control in the rescue experiment. The overexpression of *Os*PRT6-136H, *Os*PRT6-161V or double 136H+161V mutants defective in the binding of Arg-CysO_2_ peptide mimic show a significant submergence-resistant phenotype similar to the *prt6-5* lines, suggesting loss of PRT6 function in those mutants.

**Figure 5.**
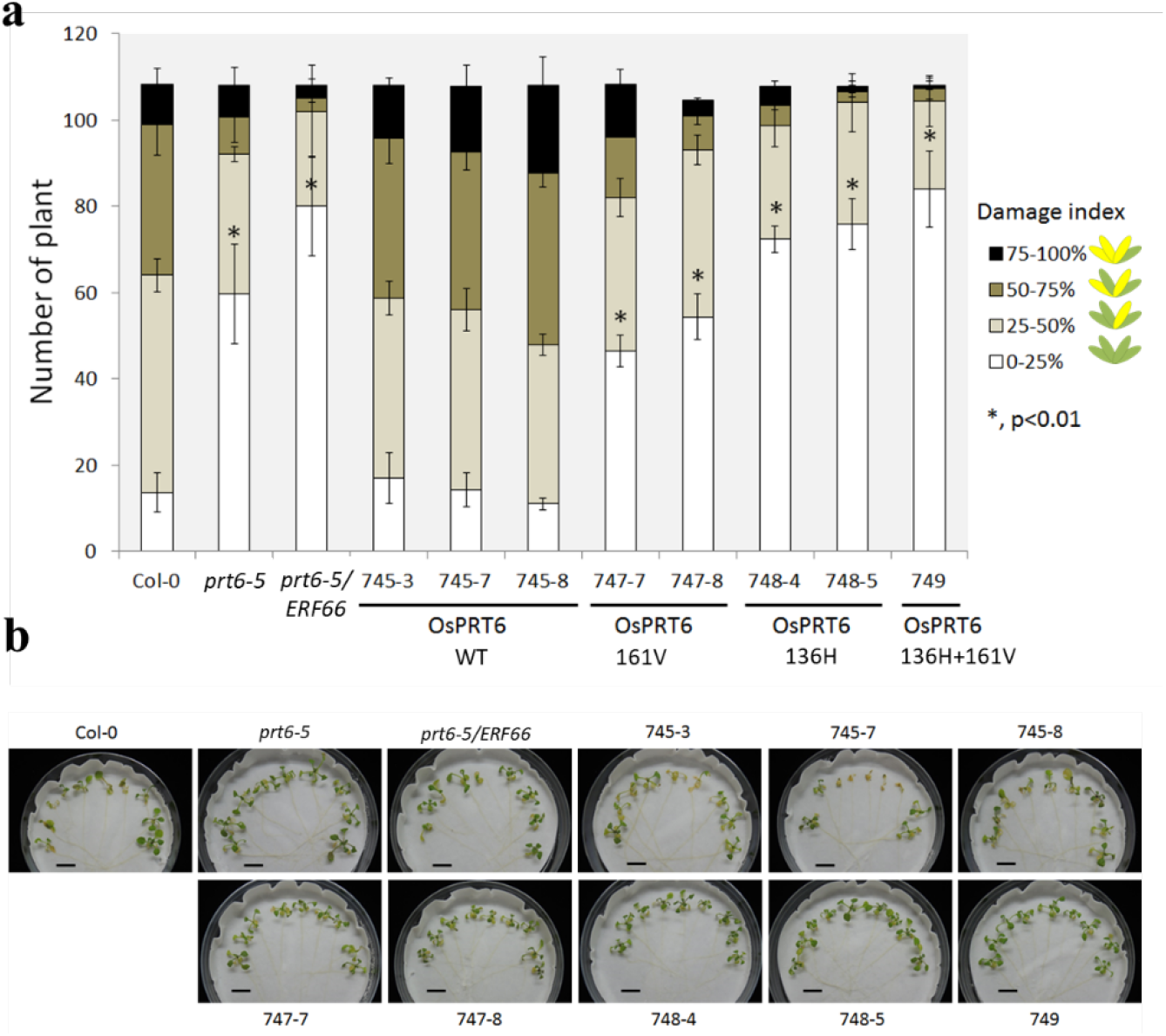
The damage index of Col-0 and *prt6-5* lines with overexpressed *Os*PRT6 variants after submergence and recovery. (a) Bar graph summarizing the damage index score of over 100 seedlings for three repeats. (b) Representatives of each line after recovery placed in Petri dishes for index score examination. After submergence and recovery, the number of chlorotic leaves over total leaves were counted for each plant as the damage index. The data represent means ± SD from three independent replicates. Statistical differences between Col-0 and *prt6-5* with overexpressed *Os*PRT6 variants were determined by Student’s *t* test. *, *P* < 0.01.

To further examine whether the degradation of the Cys-Arg/N-degron substrate is reduced in the mutant lines, an *ex vivo* protein stability assay using protoplasts was established (Fig. 5a). RT-qPCR analyses confirmed that HA-tagged *prt6* was transcribed in our overexpressed lines (Supplementary Fig. 3). Using protoplast transient assays, we overexpressed luciferase with the Cys-Arg/N-degron in PRT6-WT and PRT6-double mutant overexpression lines with similar overexpression levels of PRT6. Cycloheximide chase experiments showed that luciferase with the Cys-Arg/N-degron was degraded faster in the PRT6-WT overexpression line compared to the PRT6-double mutant overexpression line (Fig. 6 and Supplementary Fig. 4). More quantitatively, approximately 30% of the model Cys-Arg/N-degron substrate was degraded in the wild-type protoplast line within 10 min, whereas less than 10% of the substrate was degraded in the mutant line. Taken together, our phenotype and *ex vivo* data suggest that the highly conserved motifs in PRT6-UBR have critical roles in the Cys-Arg/N-degron pathway and the corresponding mutants can manipulate Cys-Arg/N-degron activities, resulting in enhanced plant submergence resistance.

**Figure 6.**
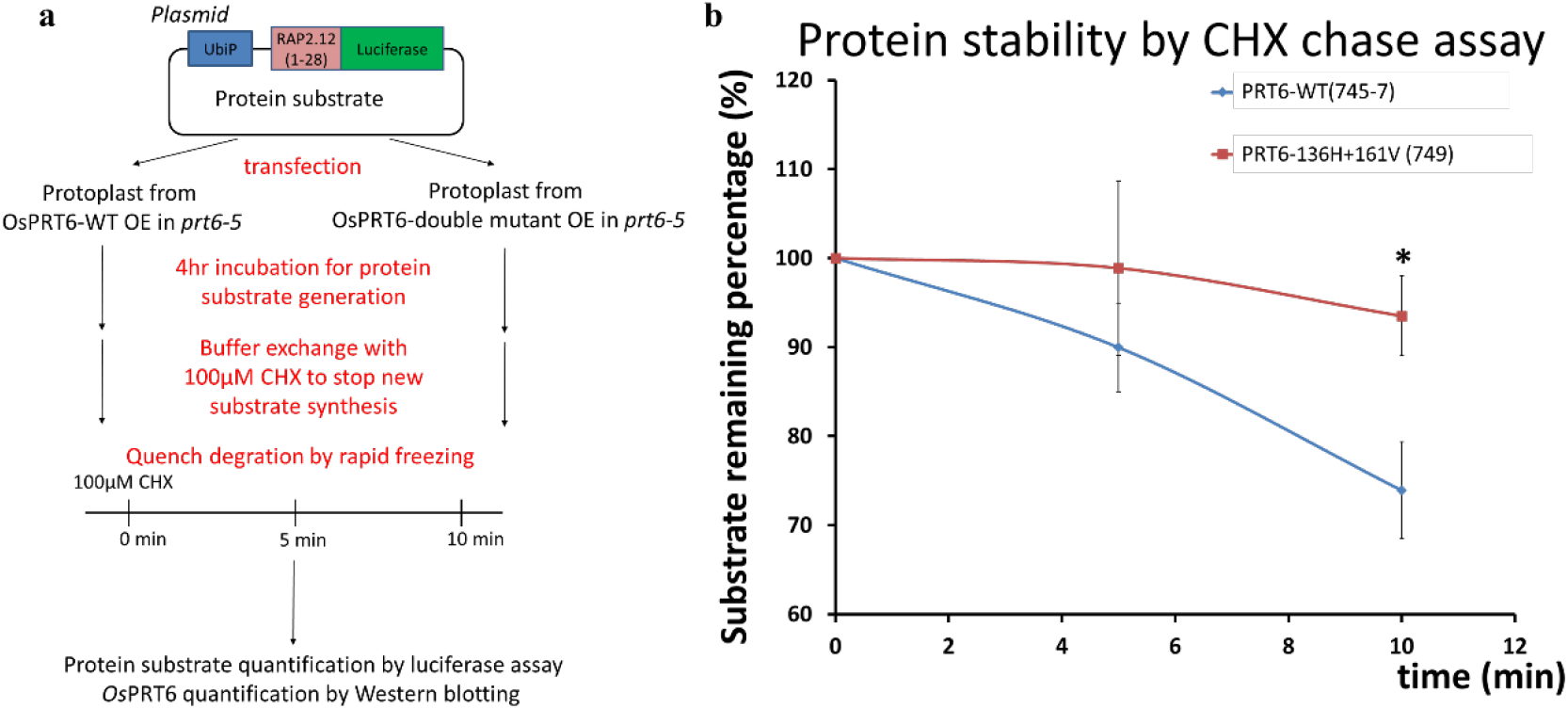
*Ex vivo* protein stability CHX chase assay. (a) Schematic procedure for *ex vivo* assay. The plasmid contained the model protein substrate, luciferase bearing Cys-Arg/N-degren sequence derived from the first 28 amino acids of *At*RAP2.12 which is a well-known ERF-VII in *Arabidopsis*. (b) The assay shows luciferase with Cys-Arg/N-degron degrades faster in the PRT6-WT overexpression line (blue) than the PRT6-double mutant (136H+161V) overexpression line (red). The data represent means ± SD from four independent replicates. Statistical differences between PRT6-WT and PRT6-double mutnat were determined by Student’s *t* test. *, *P* < 0.01.

### Concluding Remarks

This work reveals that our two newly-identified motifs in PRT6-UBR that are conserved only in plants are responsible for the Cys-Arg/N-degron. The oxygen-dependent Cys-Arg/N-degron pathway specifically generates a negatively charged cysteine sulfinic acid at the second position after the primary Arg, and is functionally critical for oxygen level sensing and responses in flowering plants^11,36^. In animals and yeast, oxygen levels are sensed by the proline hydroxylation and heme generation, respectively^37,38^. These functional differences show the uniqueness of the Cys-Arg/N-degron secondary destabilizing residue in the plant Arg/N-degron pathway. These conserved motifs can be rationalized by the requirement of the plant PRT6 to recognize the oxidized N-degron, N-terminal Arg-CysO_2_, and be evolution of the plant PRT6-UBR to enable submergence sensing. Our work thus provides important new insights into key residues with crticial roles in N-terminal Arg-CysO_2_ peptide binding in plant PRT6, and reveals further structural and mechanistic insight into how PRT6 interacts with the oxidized N-degron substrate.

Given that plant PRT6 regulates ERF-VIIs levels, and the importance of ERF-VIIs in adapting to submergence, manipulation of PRT6 activity may be an effective strategy to enhance flood tolerance in plants as shown in plant model systems^11,15,17,39^. The ability to manipulate PRT6 is important because the loss of PRT6 activity alters ABA sensitity and causes germination defects in flowering plants^15,18–20^, as shown in similar concept on PCO manipulation^40,41^. However, PRT6 is essential in the plant Arg/N-degron pathway whereas PCO contains five isoforms, which makes PRT6 a more feasible target. In this study, we demonstrate that two highly conserved regions of the plant PRT6-UBR box evolved for CysO_2_ recognition. Mutations in these regions reduce PRT6 function *in vtro* and plants with mutated PRT6 show a better submergence resistance phenotype. This structural work lays the foundations for structure-guided manipulation of PRT6 to balance benefit and harm. As climate change is increasing, the likelihood of submergence that threatens food security is increases. This work underscores the feasibility of altering PRT6 can to improve plant submergence resistance, with applications in agriculture.

## Materials and Methods

### Phylogenetic analysis

Phylogenetic analysis was conducted using MEGA 11 software. The phylogenetic tree was built using the neighbor-joining method, inferred from 1000 bootstrap replicates^42^. The following PRT6/UBR1 were compared: *Gm*PRT6 (GM07G15840, *Glycine max*), *Eg*PRT6 (EG0001G10280, *Eucalyptus grandis*), *Cs*PRT6 (CS00454G00020, *Citrus sinensis*), *Sl*PRT6 (SL10G084760, *Solanum lycopersicum*), *St*PRT6 (ST10G024170, *Solanum tuberosum*), *Pt*PRT6 (PT06G08570, *Populus trichocarpa*), *Md*PRT6 (MD04G009580, *Malus domestica*), *Egu*PRT6 (EGU2028G0858, *Elaeis guineensis*), *Pequ*PRT6 (PEQU_18062, *Phalaenopsis equestris*), *Os*PRT6 (LOC_Os01G05500, *Oryza sativa), Ph*PRT6 (PH1004749G0010, *Phyllostachys edulis), BradiPRT6* (Bradi2G03180, *Brachypodium distachyon*), *Tae*PRT6 (TAE37857G001, *Triticum aestivum), At*PRT6 (AT5G02310, *Arabidopsis thaliana), Dr*UBR1 (XP_021323192.1, *Danio rerio*), *Bp*UBR1 (XP_020775514.1, *Boleophthalmus pectinirostris*), *Cc*UBR1 (XP_023783982.1, *Cyanistes caeruleus*), *Pt*UBR1 (XP_026565212.1, *Pseusonaja textillis*), *Hs*UBR1 (NP_777576.1, *Homo sapiens*), *Mm*UBR1 (NP_033487.2, *Mus musculus*), *Rn*UBR1 (NP_001171543.1, *Rattus norvegicus*), *Sc*UBR1 (NP_011700.1, *Saccharomyces cerevisiae*). Structure alignment was calculated by PROMALS3D using *Hs*UBR1-UBR (PDB ID: 3NY1), *Hs*UBR2-UBR, (PDB ID: 3NY2), *Sc*UBR1-UBR (PDB ID: 3NIL), *Os*PRT6-UBR and *At*PRT6-UBR as inputs^43^.

### Cloning and sample production

The UBR domain of PRT6 from *Oryza sativa indica* (residues Ala120–Gly190) was cloned into a pET 15b vector by Ligation Independent Cloning (LIC) to construct a N-terminal fusion tag containing a Nt-hexahistidine, glutathione S-transferase and tobacco etch virus (TEV) protease cutting site, which leaves an additional serine residue after TEV protease treatment. The plasmid containing the gene *Os*PRT6-UBR was transformed into *E. coli* BL21 (DE3) for recombinant protein production. The transformed cells were cultured in a rich LB medium at 37°C in the presence of 100 μg/ml ampicillin and were induced at the optical density (O.D.) of 1.0 by 0.5 mM isopropyl-β-d-thiogalactoside (IPTG) at 16°C for 16 h. The harvested cells were collected by centrifugation and resuspended in lysis buffer (50 mM Tris-HCl at pH 8.0, 250 mM NaCl, 25 mM imidazole, 10% (w/v) glycerol and 10 mM β-mercatpoethanol) supplemented 1 mM PMSF and DNase followed by sonication. After centrifugation at 20,000*g* for 30 min, the supernatant was loaded onto a column containing NiNTA resin pre-equilibrated with lysis buffer. The column was washed with lysis buffer followed by 5 column volumes (CVs) of wash buffer supplemented with an additional 25 mM imidazole. Proteins were eluted with elution buffer (50 mM Tris-HCl at pH 8.0, 250 mM NaCl and 500 mM imidazole). After removing the N-terminal fusion tag by treating with TEV protease during the dialysis against 50 mM Tris buffer at pH 5.0 and 250 mM NaCl overnight at 4°C, *Os*PRT6-UBR was loaded onto a Superdex 75 column pre-equilibrated with 20 mM Tris at pH 7.5, 10 mM TCEP, 10 mM NaCl and 10 μM ZnCl_2_ for final purification.

The UBR box domain of PRT6 from *Arabidopsis thaliana* (residues Gly119-Gly189) was cloned into KpnI and XhoI restriction sites of a modified pET-His-LC3B vector^44^. For crystallization, we inserted N-degron sequences (RRGSGG and RLGSGG) between the LC3B sequence and the *At*PRT6 UBR box sequence. The plasmid containing the *At*PRT6-UBR box gene was transformed into *E. coli* BL21(DE3) cells for recombinant protein production. The plasmid-transformed *E. coli* BL21(DE3) cells were incubated in LB medium at 37°C in the presence of 100 μg/ml kanamycin until O.D. of 2.0 and then induced with a final concentration of 0.5 mM IPTG at 18°C with 500 μM ZnSO_4_ for 20 h. The harvested cells were collected by centrifugation and resuspended in a lysis buffer containing 50 mM Tris pH 8.0, 200 mM NaCl, and 1 mM Tris(2-carboxyethyl)phosphine (TCEP). Cell lysis was performed using ultrasonication supplemented with 1 mM PMSF and a tablet of protease inhibitor cocktail. The insoluble fraction was removed by centrifugation at 17,000 rpm for 2 h, and the supernatant was loaded onto a pre-equilibrated HisTrap^TM^ column (GE Healthcare, 17-5255-01) and eluted by gradually increasing the concentration of imidazole to 500 mM. Eluted samples were further purified using the HiTrap™ Q HP column (GE Healthcare, 17-1154-01). The His-LC3B-tag was cleaved using human ATG4B protease at 25°C overnight^45^. Samples with 30 mM imidazole were loaded onto the HisTrap™ column to remove the His-LC3B-tag and human ATG4B protease. Loading through samples were concentrated by ultrafiltration (Amicon Ultra 3K NMWL, Millipore) and loaded onto a HiLoad™ 16/600 Superdex™ 75 pg (GE Healthcare, 28-9893-33) column equilibrated with 20 mM Tris-HCl pH 8.0, 100 mM NaCl, and 1 mM TCEP.

For fluorescence polarization assays (FP-assays), the same construct of the UBR box domain of *Os*PRT6 was cloned into His-MBP-vectors. The HisMBP-*Os*PRT6 UBR box plasmids were transformed into *E. coli* BL21(DE3) cells. Cell lysates were applied to a column containing amylose resin (New England Biolabs). The beads were washed with at least ten column volumes of buffer containing 50 mM Tris-HCl pH 8.0, 100 mM NaCl, and 1 mM TCEP, and the MBP-*Os*PRT6 UBR box was eluted with the same buffer supplemented with 10 mM maltose. Eluted samples were further purified using the HiTrap^TM^ Q HP column, concentrated by ultrafiltration (Amicon Ultra 10K NMWL, Millipore), and then loaded onto a HiLoad^TM^ 16/600 Superdex^TM^ 200 pg (GE Healthcare, 28-9893-35) gel filtration column pre-equilibrated with 50 mM HEPES pH 7.0, 10 mM NaCl, and 1 mM TCEP.

### Protein crystallization and structure determination

*Os*PRT6-UBR was concentrated to 8–9 mg/ml prior to crystallization. Crystals were grown at 18°C by the sitting drop vapor diffusion method. Apo-form protein was crystallized in 8% (v/v) Tacsimate pH 6.0 and 20% PEG3350. An additional 20% (w/v) glycerol was added to crystallization drops and crystals were flash-cooled in liquid nitrogen at 100 K prior to data collection. For RDG/RSG peptide soaking, apo *Os*PRT6-UBR was crystallized in 1.0 M sodium citrate, 200 mM NaCl and 0.1 M Tris-HCl at pH 7.0. Crystals were transferred to 300 mM citrate/glycine/HEPES buffer at the desired pH, 1.0 M sodium citrate and 200 mM NaCl with saturated peptide for 1 h followed by flash-freezing in liquid nitrogen with an additional 25%(w/v) glycerol as a cryoprotectant. X-ray diffraction data were collected at beamline BL13B1 and TPS05A at the National Synchrotron Radiation Research Center (NSRRC, Hsinchu, Taiwan). Data were processed using the HKL2000 program suite^46^. Data processing and refinement statistics are summarized in Supplementary Table 1. Apo-form structures were determined by zinc phasing using Phenix and the peptide-bound structure was determined by molecular replacement using the apo structure as the search model^47,48^. Models with peptides were iteratively rebuilt using Coot and refined with using REFMAC5^49,50^. The bound-peptides were built only after the R-free value decreased below 30% and were guided by Fo–Fc electron density maps contoured at 3σ.

*At*PRT6-UBR box was concentrated to 7–15 mg/ml prior to crystallization. Crystals were grown at 22°C by the sitting drop vapor diffusion method. RRGSGG-*At*PRT6 UBR box (*P* 2 21 21, PDB ID: 7XWE) was crystallized in 0.2 M MgCl_2_ hexahydrate, 0.1 M sodium HEPES pH 7.5, and 30% (v/v) PEG 400; RRGSGG-*At*PRT6 UBR box (*I* 2 2 2, PDB ID: 7Y6W) in 0.2 M calcium acetate hydrate and 20% (w/v) PEG 3,350, and RRGSGG-*At*PRT6 UBR box (*P* 32, PDB ID: 7Y6X) in 1.2 M DL-malic acid pH 7.0 and 0.1 M BIS-Tris pH 7.0. RLGSGG-*At*PRT6 UBR box (*I* 2 2 2, PDB ID: 7XWF) was crystallized in 0.2 M calcium acetate hydrate and 17% (w/v) PEG 3,350; RLGSGG-*At*PRT6 UBR box (*I* 2 2 2, PDB ID: and 7Y6Z) was crystallized in 0.2 M calcium acetate hydrate and 19% (w/v) PEG 3,350; RLGSGG-*At*PRT6 UBR box (*C* 1 2 1, PDB ID: 7Y6Y) in 0.2 M calcium acetate hydrate and 20% (w/v) PEG 3,350, and RLGSGG-*At*PRT6 UBR box (*P* 4_3_ 2, PDB ID: 7Y70) in 1.3 M ammonium sulfate, 0.1 M Tris pH 8.5, and 12% (w/v) glycerol. Apo-AtPRT6 (PDB ID: 7XWD) was crystallized in 1.6 M ammonium sulfate, 0.1 M sodium HEPES pH 7.5, and 0.1 M NaCl. Crystals were usually cryoprotected by adding 10–25% (w/v) glycerol and then frozen in liquid nitrogen. X-ray diffraction data were collected at beamline 5C and 11C at the Pohang Accelerator Laboratory, South Korea, and BL44XU at the Spring-8, Japan. Data sets were processed using the HKL2000 program suite^46^. Statistics for the collected data are summarized in Supplementary Table 2. All structure figures were generated by PyMOL and LigPlot+^51,52^.

### Fluorescence polarization assay

Solutions of FITC-labeled RDAAK, RRAAK, RLAAK, and RSAAK peptides were prepared at 500 nM concentration in 50 mM MES pH 6.0, 10 mM NaCl, and 1 mM TCEP. Purified MBP-*Os*PRT6 UBR box WT and the respective mutants were serially diluted in the same buffer. Fluorescence measurements to detect the change in fluorescence polarization of the FITC-labeled peptide were performed in a 384-well format on a Corning black plate reader with excitation and emission wavelengths of 485 and 525 nm, respectively. GraphPad Prism 7 software was used for calculations and generation of graphs.

### Plant materials

The T-DNA insertion lines of *prt6-5* (SALK_051088) were obtained from the Arabidopsis Bioglocal Resoruce Center, Ohio State University. The *Os*ERF66 overexpression lines (*prt6-5/ERF66* lines) were generated by transforming the 35S::*Os*ERF66-GFP/pCAMBIA1302 vector into *prt6-5.The* HA-PRT6 WT, 136H, 161V, and 136H+161V overexpression lines were generated by transforming the *35S::HA-PRT6* WT, 136H, 161V, 136H+161V cloned *pEarleyGate 201* vector into *prt6-5/ERF66* transgenic plants.

### Damage index after submergence

*Arabidopsis* seeds were sterilized with 1.2% (v/v) sodium hypochlorite for 15 min and washed with sterilized water. Seeds were sown on plates with 0.57% Phytagel (Sigma-Aldrich) in half-strength Murashige and Skoog (MS) medium (Duchefa Biochemie) containing 0.5% sucrose at pH 5.7 and kept at 4°C in the dark for three days to achieve uniform germination, and then the plates were transferred to a growth chamber and grown at 22°C with a 16-h-light (81 μmol·m^-2^·S^-1^)/8-h-dark cycle for four days. To obtain 14-d-old plants for submergence treatment, 7-d-old seedlings were transplanted onto fresh plates, and then the plates were placed vertically to prevent roots from growing into the medium. The transplanted seedlings were grown in the growth chamber until they were 14-d-old. For submergence treatment of 14-d-old *Arabidopsis* seedlings, plates with plants on the surface of the medium were placed into sterilized water for 32–33 h. After that, the plates were taken out from the water and transferred back into the growth chamber for 4 days. The damage index^34^ was quantified based on the percentage of chlorotic leaves (>1/3 leaf area) out of total number of leaves for each seedling.

### *Ex vivo* protein stability assay

The cycloheximide (CHX) chase assay for monitoring *ex vivo* protein stability was conducted by using an *Arabidopsis* protoplast system. The RAP2.12(1-28)-luciferase chimeric protein was used as the substrate to indicate the activity of PRT6 variants. The *Arabidopsis* UBQ10 promoter, RAP2.12(1-28), and luciferase DNA fragments were cloned into pUC18 at the same time by using NEBuilder HiFi DNA Assembly Cloning Kit (M5520, NEB) to form the substrate plasmid, pUC18_AtUP::RAP2.12(1-28)-Luc. The *Arabidopsis* protoplast preparation and transformation were conducted according to a published protocol with minor modifications^53^. The substrate plasmids were transformed into the protoplasts prepared from PRT6-WT and PRT6-136H+161V overexpression line. After incubation at 22°C for 4 h, the transformed protoplasts were aliquoted followed by buffer exchange containing 100 μM CHX to stop the protein translation process. The 90% buffer was removed and protoplasts were flash-frozen in liquid nitrogen to quench the protein degradation process followed by luciferase activity assay (E1500, Promega) to determine the relative amount of RAP2.12(1-28)-luciferase chimera protein remaining at various time points. The protein levels of overexpressed PRT6-WT and PRT6-136H+161V were determined by Western blot using anti-HA antibody (16B12, BioLegend).

## Data availability

Atomic coordinates have been deposited in the PDB under the accession codes: 7WUK, 7WUL, 7WUM, and 7WUN for *Os*PRT6-UBR; 7XWD, 7XWE, and 7XWF for *At*PRT6-UBR

## Acknowledgments

We acknowledge the Synthesis Core Facility, Institute of Biological Chemistry, Academia Sinica for peptide synethsis. We thank the Biophysics Core Facility, funded by Academia Sinica Core Facility and Innovative Instrument Project (AS-CFII111-201). We thank the ABRC B101 core facility for technical support. We also thank the technical services provided by the Synchrotron Radiation Protein Crystallography Facility of the National Core Facility Program for Biotechnology, the Ministry of Science and Technology and the National Synchrotron Radiation Research Center, a national user facility supported by the Ministry of Science and Technology of Taiwan, ROC. We also thank the staff at the beamlines 5C and 11C at the Pohang Accelerator Laboratory in South Korea and at the beamline BL44XU, Spring-8, Japan, for assistance with collecting X-ray data. This study was supported by the National Research Foundation of Korea (NRF) grants from the Korean government (grant Nos. 2020R1A2C3008285, 2021M3A9I4030068 and 2022M3A9G8082638 to HKS; 2021R1A6A1A10045235 to LK) and by the Ministry of Science and Technology, ROC (grant No. 110-2628-B-001-026 to M.C.H.).

## Author Contributions

L.K, T.J.L., Y.C.C, W.H.L, M.K. and J.L.W. performed structure and binding related experiments. C.C.L, M.Y.C. and M.C.C performed plant related experiments. C.C.L., L.K., H.C.Y and M.C.H. wrote the manuscript. C.C.L., L.K., H.C.Y. and M.C.H. designed the experiments. H.K.S. and M.C.H. analysed data. H.K.S., M.C.S. and M. C.H. supervised the work.

## References

1 Oladosu, Y et al. Submergence Tolerance in Rice: Review of Mechanism, Breeding and, Future Prospects. Sustainability-Basel 12, 1632, doi:10.3390/su12041632 (2020).

2 Fukao, T., Barrera-Figueroa, B. E., Juntawong, P. & Pena-Castro, J. M. Submergence and Waterlogging Stress in Plants: A Review Highlighting Research Opportunities and Understudied Aspects. Front Plant Sci 10, 340, doi:10.3389/fpls.2019.00340 (2019).

3 Gibbs, D. J. et al. Homeostatic response to hypoxia is regulated by the N-end rule pathway in plants. Nature 479, 415–U172, doi:10.1038/nature10534 (2011).

4 Licausi, F. et al. Oxygen sensing in plants is mediated by an N-end rule pathway for protein destabilization. Nature 479, 419–U177, doi:10.1038/nature10536 (2011).

5 Bui, L. T., Giuntoli, B., Kosmacz, M., Parlanti, S. & Licausi, F. Constitutively expressed ERF-VII transcription factors redundantly activate the core anaerobic response in Arabidopsis thaliana. Plant Sci 236, 37–43, doi:10.1016/j.plantsci.2015.03.008 (2015).

6 Gibbs, D. J. et al. Group VII Ethylene Response Factors Coordinate Oxygen and Nitric Oxide Signal Transduction and Stress Responses in Plants. Plant Physiol 169, 23–31, doi:10.1104/pp.15.00338 (2015).

7 Hinz, M. et al. Arabidopsis RAP2.2: an ethylene response transcription factor that is important for hypoxia survival. Plant Physiol 153, 757–772, doi:10.1104/pp.110.155077 (2010).

8 Sasidharan, R. et al. Signal Dynamics and Interactions during Flooding Stress. Plant Physiol 176, 1106–1117, doi:10.1104/pp.17.01232 (2018).

9 Voesenek, L. et al. Submergence-Induced Ethylene Synthesis, Entrapment, and Growth in Two Plant Species with Contrasting Flooding Resistances. Plant Physiol 103, 783–791, doi:10.1104/pp.103.3.783 (1993).

10 Yang, C. Y., Hsu, F. C., Li, J. P., Wang, N. N. & Shih, M. C. The AP2/ERF transcription factor AtERF73/HRE1 modulates ethylene responses during hypoxia in Arabidopsis. Plant Physiol 156, 202–212, doi:10.1104/pp.111.172486 (2011).

11 Vicente, J. et al. The Cys-Arg/N-End Rule Pathway Is a General Sensor of Abiotic Stress in Flowering Plants. Curr Biol 27, 3183-+, doi:10.1016/j.cub.2017.09.006 (2017).

12 Varshavsky, A. N-degron and C-degron pathways of protein degradation. Proc Natl Acad Sci U S A 116, 358–366, doi:10.1073/pnas.1816596116 (2019).

13 Kwon, Y. T. et al. An essential role of N-terminal arginylation in cardiovascular development. Science 297, 96–99, doi:DOI 10.1126/science.1069531 (2002).

14 Varshavsky, A. The N-end rule pathway and regulation by proteolysis. Protein Sci 20, 1298–1345, doi:10.1002/pro.666 (2011).

15 Dissmeyer, N. Conditional Protein Function via N-Degron Pathway-Mediated Proteostasis in Stress Physiology. Annu Rev Plant Biol 70, 83–117, doi:10.1146/annurev-arplant-050718-095937 (2019).

16 Weits, D. A. et al. Plant cysteine oxidases control the oxygen-dependent branch of the N-end-rule pathway. Nature Communications 5, 3425, doi:10.1038/ncomms4425 (2014).

17 Riber, W. et al. The Greening after Extended Darkness1 Is an N-End Rule Pathway Mutant with High Tolerance to Submergence and Starvation. Plant Physiology 167, 1616–1629, doi:10.1104/pp.114.253088 (2015).

18 Graciet, E. et al. The N-end rule pathway controls multiple functions during Arabidopsis shoot and leaf development. P Natl Acad Sci USA 106, 13618–13623, doi:10.1073/pnas.0906404106 (2009).

19 Holman, T. J. et al. The N-end rule pathway promotes seed germination and establishment through removal of ABA sensitivity in Arabidopsis. P Natl Acad Sci USA 106, 4549–4554, doi:10.1073/pnas.0810280106 (2009).

20 Zhang, H. T. et al. N-terminomics reveals control of Arabidopsis seed storage proteins and proteases by the Arg/N-end rule pathway. New Phytol 218, 1106–1126, doi:10.1111/nph.14909 (2018).

21 Tasaki, T. et al. A family of mammalian E3 ubiquitin ligases that contain the UBR box motif and recognize N-degrons. Mol Cell Biol 25, 7120–7136, doi:10.1128/Mcb.25.16.7120-7136.2005 (2005).

22 Kaur, G. & Subramanian, S. The UBR-box and its relationship to binuclear RING-like treble clef zinc fingers. Biol Direct 10, 36, doi:10.1186/s13062-015-0066-5 (2015).

23 Choi, W. S. et al. Structural basis for the recognition of N-end rule substrates by the UBR box of ubiquitin ligases. Nat Struct Mol Biol 17, 1175-+, doi:10.1038/nsmb.1907 (2010).

24 Matta-Camacho, E., Kozlov, G., Li, F. F. & Gehring, K. Structural basis of substrate recognition and specificity in the N-end rule pathway. Nat Struct Mol Biol 17, 1182-+, doi:10.1038/nsmb.1894 (2010).

25 Permyakov, S. E. et al. Oxidation mimicking substitution of conservative cysteine in recoverin suppresses its membrane association. Amino Acids 42, 1435–1442, doi:10.1007/s00726-011-0843-0 (2012).

26 Urmey, A. R. & Zondlo, N. J. Cysteine oxidation to the sulfinic acid induces oxoform-specific lanthanide binding and fluorescence in a designed peptide. Free Radical Bio Med 152, 166–174 (2020).

27 Demes, E. et al. Dynamic measurement of cytosolic pH and [NO3 (-)] uncovers the role of the vacuolar transporter AtCLCa in cytosolic pH homeostasis. Proc Natl Acad Sci U S A 117, 15343–15353, doi:10.1073/pnas.2007580117 (2020).

28 Kulichikhin, K. Y., Chirkova, T. V. & Fagerstedt, K. V. Intracellular pH in rice and wheat root tips under hypoxic and anoxic conditions. Plant Signal Behav 3, 240–242, doi:10.4161/psb.3.4.5151 (2008).

29 Yemelyanov, V. V., Chirkova, T. V., Shishova, M. F. & Lindberg, S. M. Potassium Efflux and Cytosol Acidification as Primary Anoxia-Induced Events in Wheat and Rice Seedlings. Plants (Basel) 9, doi:10.3390/plants9091216 (2020).

30 Munoz-Escobar, J., Matta-Camacho, E., Cho, C., Kozlov, G. & Gehring, K. Bound Waters Mediate Binding of Diverse Substrates to a Ubiquitin Ligase. Structure 25, 719–729 e713, doi:10.1016/j.str.2017.03.004 (2017).

31 Zhou, H. X. & Pang, X. D. Electrostatic Interactions in Protein Structure, Folding, Binding, and Condensation. Chem Rev 118, 1691–1741, doi:10.1021/acs.chemrev.7b00305 (2018).

32 Ali, S. T., Karamat, S., Kona, J. & Fabian, W. M. Theoretical prediction of pKa values of seleninic, selenenic, sulfinic, and carboxylic acids by quantum-chemical methods. J Phys Chem A 114, 12470–12478, doi:10.1021/jp102266v (2010).

33 Graciet, E. et al. The N-end rule pathway controls multiple functions during Arabidopsis shoot and leaf development. Proc Natl Acad Sci U S A 106, 13618–13623, doi:10.1073/pnas.0906404106 (2009).

34 Hsu, F. C. et al. Submergence confers immunity mediated by the WRKY22 transcription factor in Arabidopsis. Plant Cell 25, 2699–2713, doi:10.1105/tpc.113.114447 (2013).

35 Lin, C. C. et al. Regulatory cascade involving transcriptional and N-end rule pathways in rice under submergence. Proc Natl Acad Sci U S A 116, 3300–3309, doi:10.1073/pnas.1818507116 (2019).

36 Abbas, M. et al. An oxygen-sensing mechanism for angiosperm adaptation to altitude. Nature 606, 565–569, doi:10.1038/s41586-022-04740-y (2022).

37 Hon, W. C. et al. Structural basis for the recognition of hydroxyproline in HIF-1 alpha by pVHL. Nature 417, 975–978, doi:10.1038/nature00767 (2002).

38 Elkins, J. M. et al. Structure of factor-inhibiting hypoxia-inducible factor (HIF) reveals mechanism of oxidative modification of HIF-1 alpha. J Biol Chem 278, 1802–1806, doi:10.1074/jbc.C200644200 (2003).

39 Mendiondo, G. M. et al. Enhanced waterlogging tolerance in barley by manipulation of expression of the N-end rule pathway E3 ligase PROTEOLYSIS6. Plant Biotechnol J 14, 40–50, doi:10.1111/pbi.12334 (2016).

40 White, M. D. et al. Structures of Arabidopsis thaliana oxygen-sensing plant cysteine oxidases 4 and 5 enable targeted manipulation of their activity. P Natl Acad Sci USA 117, 23140–23147, doi:10.1073/pnas.2000206117 (2020).

41 Taylor-Kearney, L. J. & Flashman, E. Targeting plant cysteine oxidase activity for improved submergence tolerance. Plant J, doi:10.1111/tpj.15605 (2021).

42 Tamura, K., Stecher, G. & Kumar, S. MEGA11: Molecular Evolutionary Genetics Analysis Version 11. Mol Biol Evol 38, 3022–3027, doi:10.1093/molbev/msab120 (2021).

43 Pei, J. M., Kim, B. H. & Grishin, N. V. PROMALS3D: a tool for multiple protein sequence and structure alignments. Nucleic Acids Res 36, 2295–2300, doi:10.1093/nar/gkn072 (2008).

44 Kim, L., Kwon, D. H., Heo, J., Park, M. R. & Song, H. K. Use of the LC3B-fusion technique for biochemical and structural studies of proteins involved in the N-degron pathway. J Biol Chem 295, 2590–2600, doi:10.1074/jbc.RA119.010912 (2020).

45 Kwon, D. H. et al. The 1:2 complex between RavZ and LC3 reveals a mechanism for deconjugation of LC3 on the phagophore membrane. Autophagy 13, 70–81, doi:10.1080/15548627.2016.1243199 (2017).

46 Otwinowski, Z. & Minor, W. Processing of X-ray diffraction data collected in oscillation mode. Methods Enzymol 276, 307–326 (1997).

47 Vagin, A. & Teplyakov, A. MOLREP: an automated program for molecular replacement. J Appl Crystallogr 30, 1022–1025, doi:Doi 10.1107/S0021889897006766 (1997).

48 Liebschner, D. et al. Macromolecular structure determination using X-rays, neutrons and electrons: recent developments in Phenix. Acta Crystallogr D 75, 861–877, doi:10.1107/S2059798319011471 (2019).

49 Emsley, P., Lohkamp, B., Scott, W. G. & Cowtan, K. Features and development of Coot. Acta Crystallogr D Biol Crystallogr 66, 486–501, doi:10.1107/S0907444910007493 (2010).

50 Murshudov, G. N. et al. REFMAC5 for the refinement of macromolecular crystal structures. Acta Crystallogr D Biol Crystallogr 67, 355–367, doi:10.1107/S0907444911001314 (2011).

51 Laskowski, R. A. & Swindells, M. B. LigPlot+: multiple ligand-protein interaction diagrams for drug discovery. J Chem Inf Model 51, 2778–2786, doi:10.1021/ci200227u (2011).

52 Schrodinger, LLC. The PyMOL Molecular Graphics System, Version 1.8 (2015).

53 Wu, F. H. et al. Tape-Arabidopsis Sandwich - a simpler Arabidopsis protoplast isolation method. Plant Methods 5, 16, doi:10.1186/1746-4811-5-16 (2009).

