## supplemental figures and tables for "Structural analyses of the plant PRT6-UBR box in the Cys-Arg/N-degron pathway and insights into the plant submergence response"

**Supplementary Information**

**Supplementary Figure 1. Phylogenetic tree analysis of plant PRT6-UBR, animal UBR1-UBR and *S. cerevisiae* UBR1-UBR boxes with sequence alignment.** Cysteine and histidine residues that chelate structural zinc ions are indicated as red dots and aspartate residues involved in arginine binding are indicated as blue dots. The two motifs highly conserved in the plant PRT6-UBR box are highlighted in green boxes. The residue numberings are shown on each side of the sequence alignment. The amino acid sequences of UBR boxes from the following organisms were used: *Gm*PRT6 (GM07G15840, *Glycine max*), *Eg*PRT6 (EG0001G10280, *Eucalyptus grandis*), *Cs*PRT6 (CS00454G00020, *Citrus sinensis*), *Sl*PRT6 (SL10G084760, *Solanum lycopersicum*), *St*PRT6 (ST10G024170, *Solanum tuberosum*), *Pt*PRT6 (PT06G08570, *Populus trichocarpa*), *Md*PRT6 (MD04G009580, *Malus domestica*), *Egu*PRT6 (EGU2028G0858, *Elaeis guineensis*), *Pequ*PRT6 (PEQU_18062, *Phalaenopsis equestris*), *Os*PRT6 (LOC_Os01G05500, *Oryza sativa*), *Ph*PRT6 (PH1004749G0010, *Phyllostachys edulis*), *Bradi*PRT6 (Bradi2G03180, *Brachypodium distachyon*), *Tae*PRT6 (TAE37857G001, *Triticum aestivum*), *At*PRT6 (AT5G02310, *Arabidopsis thaliana*), *Dr*UBR1 (XP_021323192.1, *Danio rerio*), *Bp*UBR1 (XP_020775514.1, *Boleophthalmus pectinirostris*), *Cc*UBR1 (XP_023783982.1, *Cyanistes caeruleus*), *Pt*UBR1 (XP_026565212.1, *Pseusonaja textillis*), *Hs*UBR1 (NP_777576.1, *Homo sapiens*), *Mm*UBR1 (NP_033487.2, *Mus musculus*), *Rn*UBR1 (NP_001171543.1, *Rattus norvegicus*), *Sc*UBR1 (NP_011700.1, *Saccharomyces cerevisiae*).

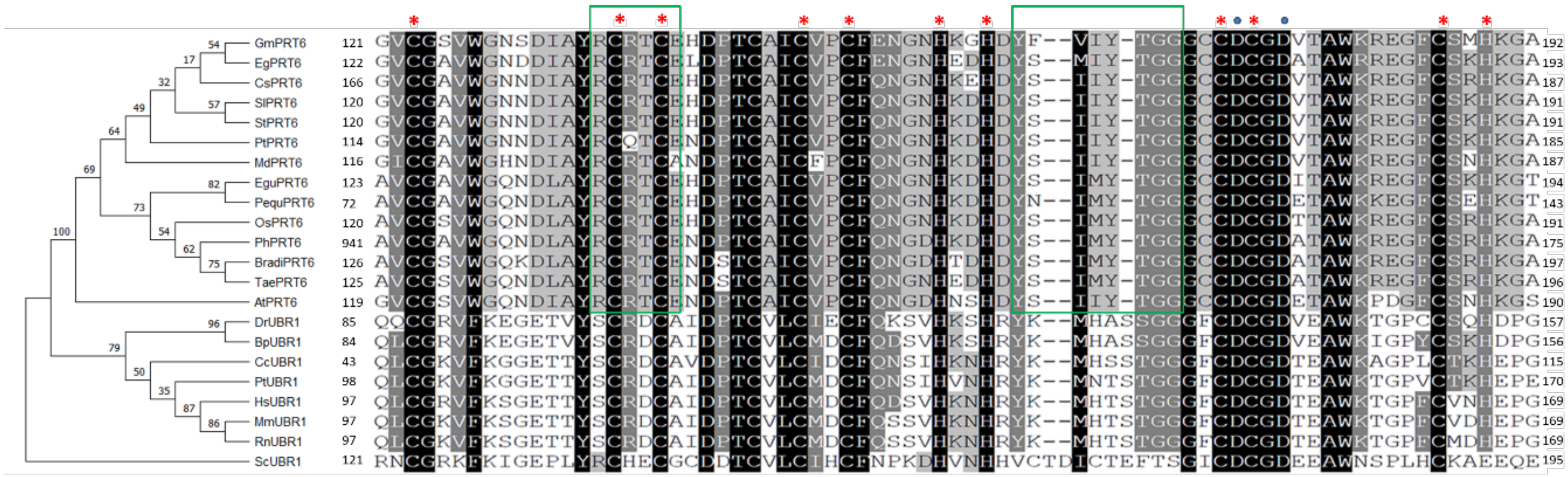

GmPRT6 121 192
EgPRT6 122 193
CsPRT6 166 187
SlPRT6 120 191
StPRT6 120 191
PtPRT6 114 185
MdPRT6 116 187
EguPRT6 123 194
PequPRT6 72 143
OsPRT6 120 191
PhPRT6 941 175
BradiPRT6 126 197
TaePRT6 125 196
AtPRT6 119 190
DrUBR1 85 157
BpUBR1 84 156
CcUBR1 43 115
PtUBR1 98 170
HsUBR1 97 169
MmUBR1 97 169
RnUBR1 97 169
ScUBR1 121 195

**Supplementary Figure 2. Structures of Arg residues of *At*PRT6-UBR with various N-degron peptides.** (a) Structure of the *At*PRT-UBR/RRG complex. (b) Structure of the *At*PRT-UBR/RLG complex. The RRG-bound and RLG-bound *At*PRT6-UBR are shown as yellow-green and brown sticks, respectively. The RRG and RLG peptide are shown as gray and magenta sticks, respectively. Water molecules are shown as red balls. The hydrogen bond and charge–charge interactions are indicated as black and green dashed lines. The orange dashed line indicates key distances.

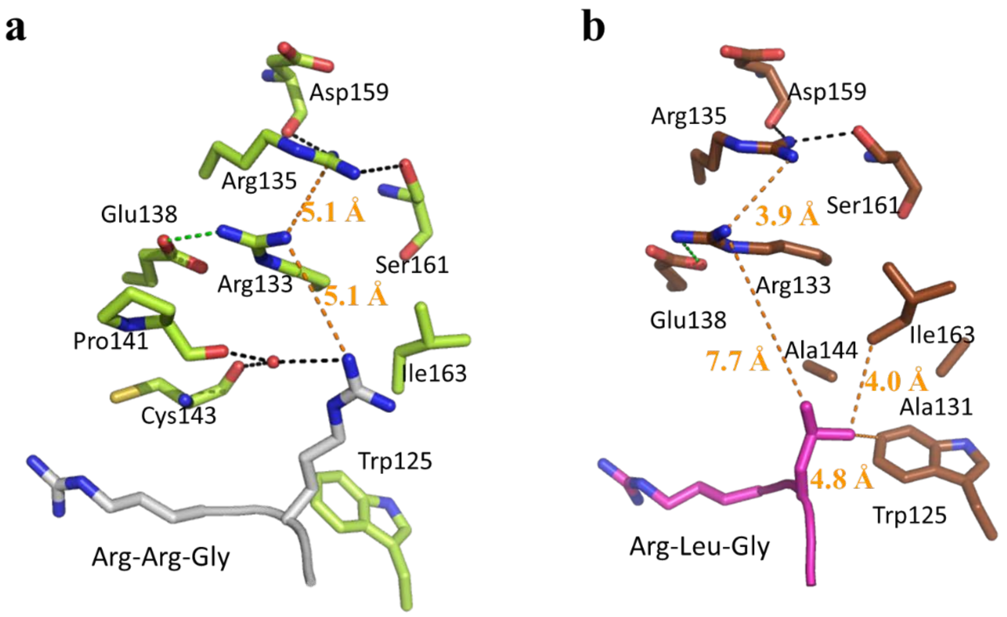

**Supplementary Figure 3. Relative mRNA levels of *Os*PRT6 variants by RT-qPCR** Tubulin (AT1G71440) was used as an internal control for normalization. Relative expression level is determined by △CT of *Os*PRT6 normalized to the internal control.

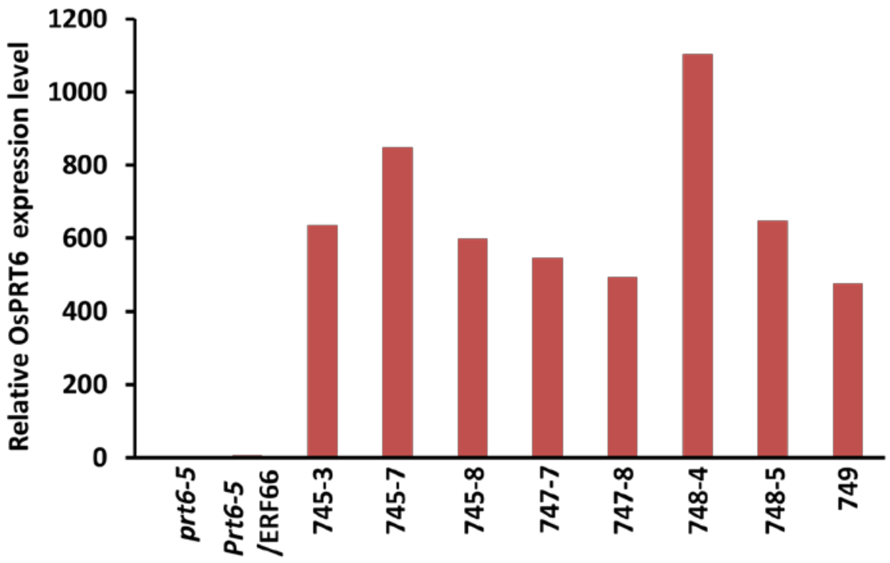


**Supplementary Figure 4. Protein levels of *Os*PRT6 variant in *ex vivo* assay by Western blot.** Lane 1: protein marker, Lane 2: *prt6-5* line protoplast, Lane 3: PRT6-WT overexpression in *prt6-5/ERF66* line protoplast, and Lane 4: PRT6-136H+161V overexpression in *prt6-5/ERF66* line protoplast. The arrow indicates the estimated molecular weight of constructed PRT6 protein in SDS-PAGE.

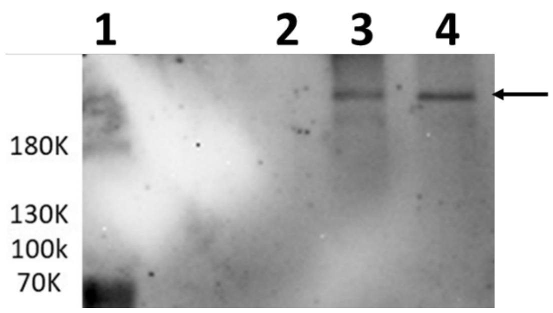

### Supplementary Table 1. Data collection and refinement statistics of *Os*PRT6-UBR box

| | **Apo** | **RDG_pH 8.5** | **RDG_pH 5.5** | **RSG_pH 5.5** |
|---|---|---|---|---|
| **PDB ID** | 7WUK | 7WUM | 7WUL | 7WUN |
| **Data Collection** | | | | |
| Wavelength (Å) | 1.2831 | 1.0 | 1.0 | 1.0 |
| Resolution range (Å) | 25.78 – 1.63 (1.69 - 1.63) | 25.77 - 1.573 (1.63 - 1.57) | 21.4 - 1.58 (1.64 - 1.58) | 23.54 - 1.53 (1.59 - 1.53) |
| Space group | $P\ 3_2\ 2\ 1$ | $P\ 3_2\ 2\ 1$ | $P\ 3_2\ 2\ 1$ | $P\ 3_2\ 2\ 1$ |
| **Cell dimensions** | | | | |
| a, b, c (Å) | 29.75, 29.75, 115.64 | 29.76, 29.76, 115.57 | 29.80, 29.80, 114.86 | 29.80, 29.80, 114.76 |
| α, γ, β (º) | 90, 90, 120 | 90, 90, 120 | 90, 90, 120 | 90, 90, 120 |
| Mean I/sigma(I) | 49.7 (15.1) | 40.1 (3.5) | 35.3 (15.7) | 20.0 (1.4) |
| R-merge | 0.04 | 0.082 | 0.06 | 0.12 |
| Completeness (%) | 99.9 (100.00) | 98.2 (69.7) | 99.5 (99.5) | 99.2 (88.4) |
| Redundancy | 6.2 (5.9) | 7.7 (3.8) | 7.6 (8.2) | 8.6 (4.1) |
| **Refinement** | | | | |
| Resolution range (Å) | 25.78 – 1.63 | 25.77 - 1.57 | 21.4 - 1.58 | 23.54 - 1.53 |
| No. reflecitons | 7,582 | 8,723 | 8,659 | 9,482 |
| $R_{work}/R_{free}$ | 0.18/0.21 | 0.18/0.21 | 0.16/0.19 | 0.17/0.19 |
| *No. atoms* | | | | |
| macromolecules | 531 | 574 | 577 | 575 |
| ligands | 3 | 3 | 3 | 3 |
| solvent | 23 | 42 | 75 | 86 |
| *Average B-factors (Å$^2$)* | | | | |
| macromolecules | 17.05 | 28.05 | 17.83 | 16.45 |
| ligands | 19.86 | 21.89 | 12.91 | 12.97 |
| solvent | 22.11 | 35.76 | 30.02 | 27.32 |
| *R.m.s. deviations* | | | | |
| Bond length (Å) | 0.013 | 0.009 | 0.007 | 0.007 |
| Bond angles (º) | 1.69 | 1.12 | 0.96 | 0.98 |
| Ramachandran favored (%) | 100.00 | 100.00 | 100.00 | 100.00 |
| Rotamer outliers (%) | 0.00 | 1.59 | 0.00 | 0.00 |

Statistics for the highest-resolution shell are shown in parentheses.

### Supplementary Table 2. Data collection and refinement statistics of *At*PRT6-UBR box

| | Apo | RRGSGG | RLGSGG |
|---|---|---|---|
| **PDB ID** | 7XWD | 7XWE | 7XWF |
| **Data Collection** | | | |
| Wavelength (Å) | 0.9 | 1.0 | 1.0 |
| Resolution range (Å) | 43.58 – 2.396 | 50.00 – 1.60 | 50.00 – 1.45 |
| | (2.482 – 2.396) | (1.63 – 1.60) | (1.48 – 1.45) |
| Space group | $I\ 4_1\ 2\ 2$ | $P\ 2\ 2_1\ 2_1$ | $I\ 2\ 2\ 2$ |
| *Cell dimensions* | | | |
| a, b, c (Å) | 87.17, 87.17, 74.35 | 46.19, 55.86, 57.32 | 39.62, 45.80, 87.09 |
| α, γ, β (º) | 90, 90, 90 | 90, 90, 90 | 90, 90, 90 |
| I/sigma(I) | 14.06 (1.52) | 12.92 (1.62) | 28.86 (1.88) |
| R-merge | 0.04 | 0.15 | 0.08 |
| Completeness (%) | 99.88 (99.31) | 99.43 (96.31) | 90.47 (59.79) |
| Redundancy | 2.0 (2.0) | 5.9 (3.2) | 7.7 (4.8) |
| **Refinement** | | | |
| Resolution range (Å) | 43.58 – 2.40 | 40.01 – 1.60 | 36.06 – 1.45 |
| No. reflections | 5,881 | 20,132 | 13,057 |
| $R_{work}/R_{free}$ | 0.22/0.27 | 0.18/0.20 | 0.19/0.21 |
| *No. atoms* | | | |
| macromolecules | 526 | 1,130 | 548 |
| ligands | 3 | 7 | 3 |
| solvent | 0 | 38 | 20 |
| *Average B-factors (Å$^2$)* | | | |
| macromolecules | 67.04 | 18.52 | 19.55 |
| ligands | 64.03 | 14.53 | 15.03 |
| solvent | 0 | 21.70 | 20.79 |
| *R.m.s. deviations* | | | |
| Bond length (Å) | 0.041 | 0.009 | 0.007 |
| Bond angles (º) | 1.21 | 1.04 | 0.98 |
| Ramachandran favored (%) | 97.06 | 99.32 | 100.00 |
| Rotamer outliers (%) | 0.00 | 0.00 | 0.00 |

Statistics for the highest-resolution shell are shown in parentheses.
